# Test Agreement Between Roche Cobas 6800 and Cepheid GeneXpert Xpress SARS-CoV-2 Assays at High Cycle Threshold Ranges

**DOI:** 10.1101/2020.05.05.078501

**Authors:** Kari Broder, Ahmed Babiker, Charles Myers, Terri White, Heather Jones, John Cardella, Eileen M. Burd, Charles E. Hill, Colleen S. Kraft

## Abstract

The SARS-CoV-2 pandemic has changed the face of the globe and upended the daily lives of billions. In an effort to bring mass-testing to as many as possible, multiple diagnostic tests including molecular, antigen detection and serological assays have been rapidly developed. However, there is very little information on positive test agreement across modalities, especially for lower viral loads. Thirty-five nasopharyngeal samples that had cycle threshold (Ct) values greater than 30.0 from the Roche cobas 6800 assay were run on the Cepheid GeneXpert Xpress SARS-CoV-2 assay. Ct values ranged from 30.1 to 37.9 (mean 36.7 ± 1.9) on the Roche cobas 6800 assay and 24.6 to 42.4 (mean 32.8±4.1) on the Cepheid assay. There was a bias of 0.33 ± 3.21, (mean difference −1.59, 95% limits of agreement −5.97, 6.63) signifying close agreement between the 2 instruments with a high standard deviation. The close test agreement between the cobas 6800 and GeneXpert at high Ct values allows for utilization of both assays interchangeable in accordance with testing algorithms.

## Introduction

Since the initial outbreak of SARS-CoV-2 on December 12, 2019 (1), in Wuhan, China, the pandemic has infected over 2.5 million people in 185 countries and is attributed to over 186,000 deaths as of April 23^rd^ 2020 (2). Due to limitations in testing availability it is likely that these case detections do not reflect the full extent of this devastating emerging virus. As part of the tremendous response effort, multiple diagnostic tests including molecular, antigen detection and serological assays have been rapidly developed. Under Food Drug Administration (FDA) Emergency Use Authorization (EUA), several RT-PCR assays have reached US laboratories, each with its own testing capacity and proprietary methods. However, there have been mutiple challenges, ranging from regulatory hurdles, reagents backlogs, swab shortages and staffing logistics (3). This has necessitated many laboratories to validate and implement multiple platforms for testing. Clinically, there is also great concern about how to interpret the results and if the results are interchangeable when laboratories have implemented a myriad of platforms. Data on positive percent agreement across platforms is limited. Our institution utilizes the Roche cobas 6800 SARS-CoV-2 (Roche Diagnostics, Indianapolis, IN), the Cepheid GeneXpert Xpress SARS-CoV-2 assay (Cepheid Inc, Sunnyvale, CA), and a laboratory developed test (LDT) based on a modified CDC protocol. The cobas 6800 assay is a qualitative dual target assay that uses a specific ORF1 target unique to SARS-CoV-2, in addition to a region on the E gene, which is conserved across the sarbecovirus subgenus (https://www.fda.gov/media/136047/download). The GeneXpert is also a qualitative dual target assay that uses the SARS-CoV-2 specific N2 region of the N gene and a region on the pan-sarbecovirus E gene (https://www.fda.gov/media/136314/download). Since there is not a gold standard for diagnostic accuracy of these SARS-CoV-2 assays, the objective was to determine the degree of agreement between the two tests by running the same samples across both platforms and comparing the cycle threshold (Ct) values comparing E target to E target among samples with high Ct values (>30), corresponding to lower levels of nucleic acid present in the sample.

## Materials and Methods

Thirty-five nasopharyngeal samples collected between April 15 – April 22, 2020 with a low positive result (Ct value ≥ 30) on the Roche cobas 6800 assay of either target underwent secondary testing on the Cepheid GeneXpert. Discrepancies were resolved using the LDT. All specimens were tested using the manufacturer’s protocol. E target Ct values were compared by Bland-Altman analysis (4) using R version 3.6.1.

## Results

Out of 35 samples, 34 tested positive on both instruments. One sample tested positive on the cobas 6800 assay (Ct=37.9), negative by the GeneXpert assay, and was confirmed to be negative on the LDT. Among positive samples, Ct values for the E gene target were similar between the two assays (p=0.06). They ranged from 30.1 to 37.9 (mean 36.7 ± 1.88) on the Roche cobas 6800 assay and 24.6 to 42.4 (mean 32.8 ± 4.07) on the GeneXpert assay. Bland-Altman analysis revealed a bias of 0.328 ± 3.21, (mean difference −1.59, 95% limits of agreement −5.97, 6.63) signifying close agreement between the 2 instruments with a high standard deviation. (**Figure 1, Table 1**).

**Table 1.** Assay Cycle Threshold Statistics. Ct data from E target gene of SARS-CoV-2 as tested on the Roche cobas 6800 and Cepheid GeneXpert assays (n=35).

|  | **Roche cobas 6800** | **Cepheid GeneXpert** |
| --- | --- | --- |
| **Median** | 33.1 | 32.2 |
| **Mean** | 33.1 | 32.8 |
| **Standard deviation** | 2.0 | 4.1 |
| **Range** | 30.1 - 37.1 | 24.6 – 42.4 |
| **Inter-quartile range** | 2.8 | 5.7 |

**Figure 1.**
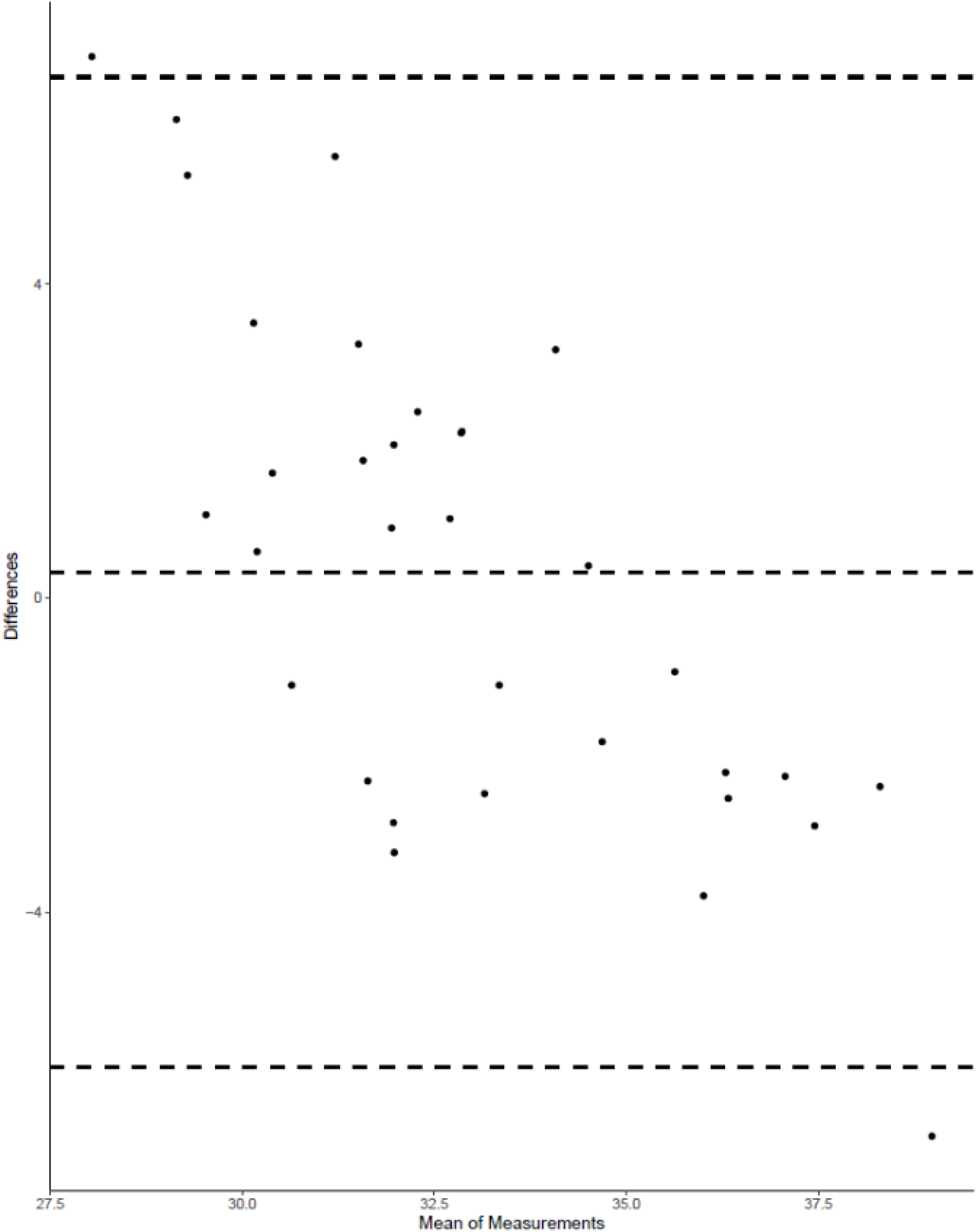
Bland Altman Plot of Agreement. Bland Altman plot showing Ct values with bias of 0.33 ± 3.21, (95% limits of agreement −5.97, 6.63).

## Discussion

Our findings corroborate recently published results which show close agreement between the cobas 6800 and GeneXpert instruments, in particular among samples with lower viral loads (5). Our in-house determined, the limit of detection (LoD) of the cobas 6800 assay is 100 copies/mL and package insert LoD on the GeneXpert. 250 copies/mL. This difference in LoD becomes significant at higher Ct values where the negative bias becomes more pronounced (4, 6). Previous studies have shown that substantial viral loads can be detected around day 5 of infection and drop at differing rates over the course of the illness (7). At our institution, the GeneXpert is utilized as a rapid test (turn-around time 45 minutes) to triage patients for admission among other criteria (**see Table 2**), therefore it is imperative that our testing modality can capture patients who present at either end of their illness course and who may have lower viral counts/higher Ct values.

**Table 2.** Healthcare algorithms for testing modality. Healthcare algorithm for the use of rapid tests, designed to maximize workflow with limited rapid test availability. * indicates rapid test being utilized Abbreviations: L&D; labor and delivery, PUI, person under investigation; SNF, skilled nursing facility

| **Patient Location** | **Additional Field** | **Detail Value** |
| --- | --- | --- |
| **ED** | Patient Status | Observation unit* |
|  |  | Inpatient |
|  |  | Outpatient |
|  | PUI Communal Dwelling | Yes* |
|  |  | No |
|  | Non-invasive Ventilation | Yes* |
|  |  | No |
| **Inpatient** | Organ Transplant Offer | Yes* |
|  |  | No |
|  | PUI Same-Day SNF or Hospice discharge | Yes* |
|  |  | No |
|  | PUI Needs ICU Care* | Yes* |
|  |  | No |
| **Labor and Delivery (L&D)** | L&D COVID Test Designator | Pregnant PUI active labor/impending delivery* |
|  |  | Non-PUI active labor/impending delivery |
|  |  | Pregnant, not impending delivery |
|  |  | Pregnant PUI being |
|  |  | discharged from Triage |
|  |  | Non-PUI being discharged<br>from Triage |

Limitations of this study include sample size, location within a single institutional system, and proprietary manufacturer information, including differences in E gene target sequence. Further studies are required to confirm percent agreement across different platforms and specimen types to expand on our findings. This is of particular importance as there is a lack of standardization of reference materials and quantitfication (copies/vol, TCID50, etc) for LoD determination. Overall, the Cepheid GeneXpert Xpress SARS-CoV-2 assay and the Roche cobas 6800 SARS-CoV-2 assay showed a high level of agreement among patients with high Ct values. This allows for the laboratories to utilize both assays concurrently as fits with local testing algorithms.

## Acknowledgements

Dr. Kraft participated on a Roche advisory board regarding COVID serology. All other authors have no conflicts. We would like to acknowledge the staff in the molecular and microbiology laboratories for their assistance in this study.

## References

1. Zhou P, Yang X-L, Wang X-G, Hu B, Zhang L, Zhang W, Si H-R, Zhu Y, Li B, Huang C-L, Chen H-D, Chen J, Luo Y, Guo H, Jiang R-D, Liu M-Q, Chen Y, Shen X-R, Wang X, Zheng X-S, Zhao K, Chen Q-J, Deng F, Liu L-L, Yan B, Zhan F-X, Wang Y-Y, Xiao G-F, Shi Z-L. 2020. A pneumonia outbreak associated with a new coronavirus of probable bat origin. Nature 579:270–273.

2. Dong E, Du H, Gardner L. 2020. An interactive web-based dashboard to track COVID-19 in real time. Lancet Infect Dis doi: 10.1016/s1473-3099(20)30120-1.

3. Babiker A, Myers CW, Hill CE, Guarner J. 2020. SARS-CoV-2 Testing: Trials and Tribulations. American Journal of Clinical Pathology doi: 10.1093/ajcp/aqaa052.

4. Giavarina D. 2015. Understanding Bland Altman analysis. Biochemia medica 25:141–151.

5. Moran A, Beavis KG, Matushek SM, Ciaglia C, Francois N, Tesic V, Love N. 2020. The Detection of SARS-CoV-2 using the Cepheid Xpert Xpress SARS-CoV-2 and Roche cobas SARS-CoV-2 Assays. Journal of Clinical Microbiology doi: 10.1128/JCM.00772-20:JCM.00772-20.

6. Ranganathan P, Pramesh CS, Aggarwal R. 2016. Common pitfalls in statistical analysis: Intention-to-treat versus per-protocol analysis. Perspectives in clinical research 7:144–146.

7. Wölfel R, Corman VM, Guggemos W, Seilmaier M, Zange S, Müller MA, Niemeyer D, Jones TC, Vollmar P, Rothe C, Hoelscher M, Bleicker T, Brünink S, Schneider J, Ehmann R, Zwirglmaier K, Drosten C, Wendtner C. 2020. Virological assessment of hospitalized patients with COVID-2019. Nature doi: 10.1038/s41586-020-2196-x.

